# Off the shelf: Investigating transfer of learning using commercially available virtual reality equipment

**DOI:** 10.1101/2022.12.19.521075

**Authors:** Logan Taylor Markwell, Koleton Cochran, Jared M. Porter

## Abstract

The evolution of virtual reality (VR) has created the opportunity for a relatively low-cost and accessible method to practice motor skills. Previous studies have demonstrated how motor skill practice in non-immersive virtual environments transfers to physical environments. Though minimal research has investigated motor learning and transfer within immersive VR, multiple experiments provide empirical evidence of positive transfer effects. To enhance the similarities between virtual and physical environments, most studies have used software engines and modified hardware. However, many learners and practitioners are currently using commercially available VR with the goal of enhancing real-world performance, though there is very little evidence to support the notion of positive transfer for these systems. Therefore, the purpose of this experiment was to investigate how motor skill practice using a commercially available VR system improved real-world performance and how that compared to physical practice. Participants (n = 68) were randomly selected into one of two groups: virtual reality (VR) practice (n = 33) or real-world (RW) practice (n = 35). The experiment took place over two consecutive days with a pretest, posttest, and practice phase occurring on both days. The pre- and post-testing phases were identical for both groups and consisted of putting a golf ball 10 times on a carpeted surface towards the center of a target. The practice phases consisted of 60 total putts per day in the respective environment (VR or RW). Participants continuously alternated golf putting holes from three different distances until they accrued 60 total putts. Participants in the RW group performed golf putts to three targets. Participants in the VR group also performed golf putts on three different miniature golf putting holes, using the commercially available Oculus Rift and the Cloudlands VR Minigolf game. The VR putting targets were designed to replicate the putting holes in the physical environment. Separate 2 (condition) x 4 (test phase) repeated measures ANOVAs were used to assess accuracy and club head kinematics. The results revealed a significant main effect for test phase, but not for condition. Post hoc analyses revealed both groups significantly improved their putting accuracy and club head kinematics at similar rates. The results from this study indicate that the transfer of learning that occurred from the commercially available VR practice was equally effective when compared to RW practice.

## Introduction

Virtual reality (VR) has recently gained popularity as a method for motor skill development. Consider a surgeon who needs to practice a specific suture, a baseball player who needs to practice hitting a knuckleball, or a pilot who needs access to a helicopter and finances to cover the cost of flying the aircraft. The evolution of immersive VR (e.g., VR systems in which the user wears a head mount display and is fully encapsulated within the virtual environment) has created the potential for a relatively low-cost method of practice that is commercially available. Within immersive VR, it is important to differentiate between animated VR, in which users can interact with the virtual environment, and 360-degree video VR, in which users are presented with a 360-degree video of prerecorded real-world video footage (1). The main difference between the two virtual environments is that users cannot interact with the environment within 360-degree video VR. A significant benefit of animated immersive VR is the opportunity for behaviors to be practiced and assessed in a challenging, yet safe and controlled environment. Furthermore, one of the largest benefits of animated VR is the ability to instantaneously adapt to the challenge level based on the individual’s skill level, maximizing learning potential (2).

Non-immersive VR (e.g., non-head mount displays) has been explored within many domains, such as surgery (3), firefighting (4), aviation (5), rehabilitation (6), and sport (2). These investigations have found a range of benefits for using VR as a form of training (2,3,4,5,6). However, there remains a scarcity of research examining how skills learned in an immersive virtual environment transfer out of the virtual space and into a physical environment. Among the studies that have investigated the effect of transfer of learning from animated VR to a physical environment (e.g., 7,8,9,10), the results have been mixed. For example, multiple experiments provided evidence that practicing a motor skill in immersive VR can increase performance within a physical environment (8 experiment 2,9,10). However, other experiments have not supported this conclusion (7,8 experiment 1). The studies above have used a variety of methods. As a result, the precise factors that contribute to transfer of learning from VR to the real world remain unclear.

Interestingly, most studies investigating this topic have used animated VR technology that utilizes customized software or software engines designed to alter the virtual environment (e.g., Unity; Unreal). For example, the Unity Experiment Framework (UXF) developed by Brookes et al. (11) has been used during human behavior research using VR (see 8 for an example). UXF allows the researcher to modify independent variables and create highly controlled testing conditions within the virtual environment that are nearly identical to the physical environment, increasing the physical and psychological similarities between the two realities (VR & real-world). In addition to customized software, the hardware is frequently modified to further enhance the physical similarities between the VR and real-world task. For example, Harris et al. (8) used a physical golf club and attached sensors to create the VR club. Similarly, Oagaz et al. (10) used a VR table tennis racket which had a similar size, weight, and shape compared to a physical racket. These software customizations and hardware modifications have provided obvious methodological benefits for testing real-world performance improvements through the use of VR. However, in a practical setting, it is unlikely that practitioners (e.g., physical therapist, coach, flight instructor) or learners (e.g., patients, athletes, students) will possess the needed skills to program a software engine (e.g., UXF) that can modify the virtual environment to simulate the physical environment with high fidelity. Likewise, not all users will have modified hardware to use during VR training. Instead, some will ultimately use “off-the-shelf” commercially available applications and hand-held controllers. The lack of physical and psychological similarities when using non-customized commercially available VR hardware and software might alter the extent to which transfer of learning occurs from a virtual to a real-world setting.

Several explanations have been proposed to explain transfer of learning. For example, the identical elements theory (12), later evolved into Singley and Andersons’s (13) identical production model, and the transfer-appropriate processing theory (14) are traditional motor learning theories that have dominated the literature. These theories propose to achieve a positive transfer of learning, similarities must exist between the practice and transfer conditions (e.g., practice specificity; 15). However, the elements theory posits that the positive transfer is due to the similarities between the movement characteristics (e.g., the swing of a golf club) and/or the environmental context in which the skill is performed (12). On the other hand, the transfer-appropriate processing theory suggests that transfer occurs due to cognitive processing similarities (14). The cognitive process similarities can be considered the type or amount of information the individual must process within the practice environment (e.g., intrinsic & extrinsic feedback). Evidence from testing the two explanations suggests there is merit to both (13, 14). Thus, it can be expected that the degree to which positive transfer of learning occurs is related to the shared similarities of the skill characteristics, environmental context, and cognitive processes. Such explanations also align with the practice specificity literature. Specifically, research testing the predictions of the practice specificity hypothesis suggest that if sources of information that were available during the learning phase of a skill are removed, performance is likely to deteriorate (15,16). Given that many software and hardware companies are marketing VR systems as a method to enhance real-world performance, understanding whether these devices can be taken off the shelf and used without modifications and what VR factors may or may not contribute to a positive transfer of learning is imperative to investigate from both a practical and theoretical perspective.

To our knowledge, only two studies have examined the transfer of learning effects from a virtual environment to a real-world environment using non-customized, commercially available immersive VR hardware and software (7,9). Michalski et al. (9) investigated how table tennis practice in immersive VR compared to a no-practice control group. Participants performed a total of three hours and 30 minutes of table tennis using a readily available application. Their analysis showed that the VR group significantly outperformed the control group, suggesting that immersive VR was beneficial for improving performance compared to no practice at all. More recently, Drew et al. (7) compared performance and kinematic differences between dart-throwing practice in immersive VR and the real world. Both groups completed 10 dart throws until they accrued a total of 100 throws and the VR group used a commercially available application and hand-held controller during the practice session. The results of Drew et al. (7) demonstrated that dart throwing accuracy significantly decreased following practice in VR while accuracy increased following real-world practice, as evidenced by a posttest in the real-world immediately following practice. However, the results also showed that there were no kinematic differences during the posttest. Though both practice groups led to similar movement characteristics during the posttest, real-world dart throwing not only outperformed VR dart throwing during the real-world posttest, but contrary to Michalski et al. (9), practice in VR led to worse real-world performance compared to the pretest. In other words, the findings reported by Drew et al. (7) demonstrated that practicing a motor skill in VR depressed the performance of the same skill when practiced in the real-world. This decrease in performance likely occurred due to the lack of similarities between conditions. This study (7) and others (1), highlight the necessity to understand how VR transfers to real-world performance.

The limited evidence examining the effects of real-world performance improvements using commercially available animated VR is mixed, and it is unclear whether VR practice is as effective as real-world practice. Specifically, to our knowledge, Drew et al. (7) is the only study that has compared VR practice, without software or hardware customizations, to real-world practice of the same task. Therefore, the purpose of the present study was to examine the transfer of learning effects using commercially available VR software and hardware during practice. Using a golf putting task, we compared real-world accuracy and club swing kinematics after VR or real-world practice. Though a VR hand-held controller was used, we predicted that both forms of practice would elicit similar accuracy improvements due to the similarities between the movement and environment characteristics as well as the cognitive processes involved. This prediction was based on our understanding of the transfer of learning effects (13,14,15) and previous studies that provided initial evidence of positive transfer (8 experiment 2,9,10). Additionally, we hypothesized that analyses of the kinematics during the posttests would reveal similarities between groups, consistent with previous research (7).

## Method

### Participants

Participants (n = 68) were recruited from undergraduate kinesiology classes to participate in this study. The participants were informed that they would practice a golf putting task but were naïve to the purpose of the study. All participants read and signed an informed consent prior to participation, and all forms and methods were approved by the university’s Institutional Review Board.

### Apparatus and task

The data collected for this experiment took place in a climate-controlled research laboratory. A golf putting task was used for both groups (VR practice; RW practice). The RW practice condition consisted of putting golf balls towards holes at three different distances (.91 m, 1.37 m, 1.83 m) on a carpeted surface inside a climate controlled research laboratory. For all pre and post-testing trials, participants used a standard length (90 cm) golf putter to putt a regular-sized (diameter 4.27 cm) golf ball towards a target. The target was a series of concentric circles. The center circle had a diameter of 10.8 cm and each concentric ring had a diameter that increased by 10.8 cm. The concentric ring in which the ball came to rest determined the score for each putt. The center circle resulted in a score of zero, a score of one was recorded if the ball came to rest in the next circle, and so on for each respective ring out to a 15^th^ circle. A score of 16 was recorded if the ball came to rest outside of the last ring. If the ball came to rest on a line of any ring, the participant received a score for the innermost ring.

The Oculus Rift VR headset and the Cloudlands VR Minigolf application were used to create three virtual golf putting holes designed to replicate the putting holes in the RW environment by shape and length. Participants used a virtual golf putter and ball to putt into a virtual hole while wearing the Oculus Rift headset and holding one Oculus controller in their dominant hand.

### Procedure

Participants were randomly assigned into one of two groups: VR practice (n = 33) or RW practice (n = 35). After participants signed the consent form, the researcher provided instructions followed by a demonstration of the golf putting task. The participant was instructed to hit the ball onto the center of the target or as close to the center of the target as possible.

The experiment took place over two consecutive days, with a pretest phase, a practice phase, and a posttest phase occurring on both days (see figure 1). The pre- and post-test phases were identical for both groups. During the pre- and post-testing phases, participants putted a golf ball 10 times on the carpeted surface toward the center of the target from a distance of 1.83 m. The practice phases consisted of 60 total putts within the respective environment (i.e., RW or VR). During the practice phases, participants continuously alternated golf putting holes from all three distances (i.e., .91 m, 1.37 m, 1.83 m) until they had accrued 60 total putts for each day, regardless of the group. The participants returned 24 hours later to complete a pretest, which also served as a delayed retention test for the RW group, or a transfer test for the VR group, from day 1, followed by a practice phase of 60 total putts and a posttest. Day one and day two were identical in structure. At the end of the second day, participants accrued 120 total putts during both practice phases.

**Figure 1.**
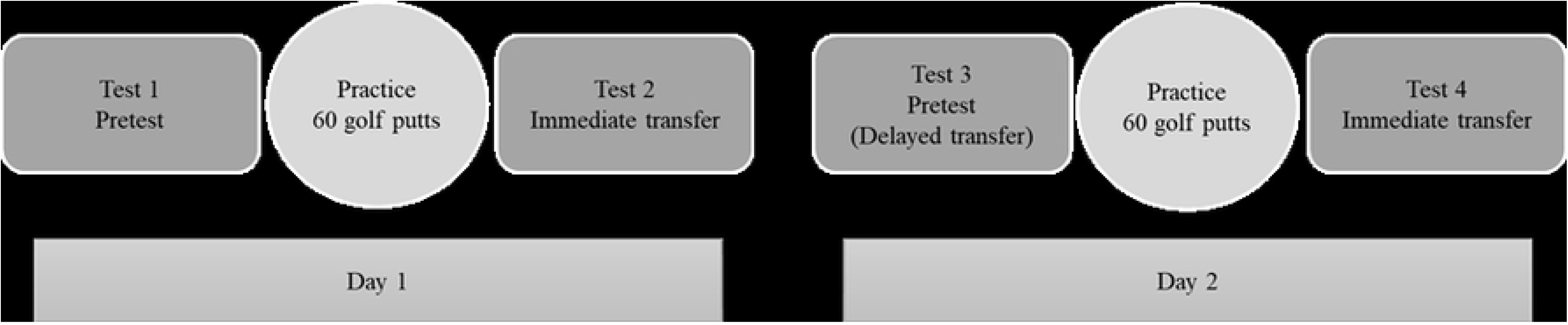
Schematic representation of the experimental procedures used during days one and two of the experiment.

### Statistical Analysis

There were four separate 2 (group) x 4 (test) repeated measures analysis of variance (ANOVAs) used to assess accuracy, backswing displacement, follow-through displacement, and club velocity differences between groups and across tests.

## Results

### Accuracy

A 2 (group) x 4 (test) repeated measures analysis of variance (ANOVA) was used to determine accuracy differences between groups and tests. Mauchly’s test indicated that the assumption of sphericity was violated (χ^2^(5) = 19.227, *p* = .002), therefore degrees of freedom were corrected using Greenhouse-Geisser estimates of sphericity (ɛ = .828). The analysis revealed a significant main effect for test *F*(2.485, 156.574) = 5.693, *p* = .002, *η_p_^2^* = .083. The interaction between test and group was not significant, *p* = .298. Furthermore, the test of between-subject effects revealed a non-significant effect, *p* = .660. In light of the significant main effect for test, pairwise comparisons were made to determine differences across tests (see figure 2). The analysis revealed that test three (*M* = 6.986, *SD* = 1.698) was significantly lower compared to test one (*M* = 7.964, *SD* = 2.436), *p* = .006. The analysis also revealed that test four (*M* = 6.857, *SD* = 2.050) was significantly lower compared to test one, *p* = .002. Additionally, the analysis revealed that test four (*M* = 6.857, *SD* = 2.050) was significantly lower compared to test two (*M* = 7.418, *SD* = 2.108), *p* = .029.

**Figure 2.**
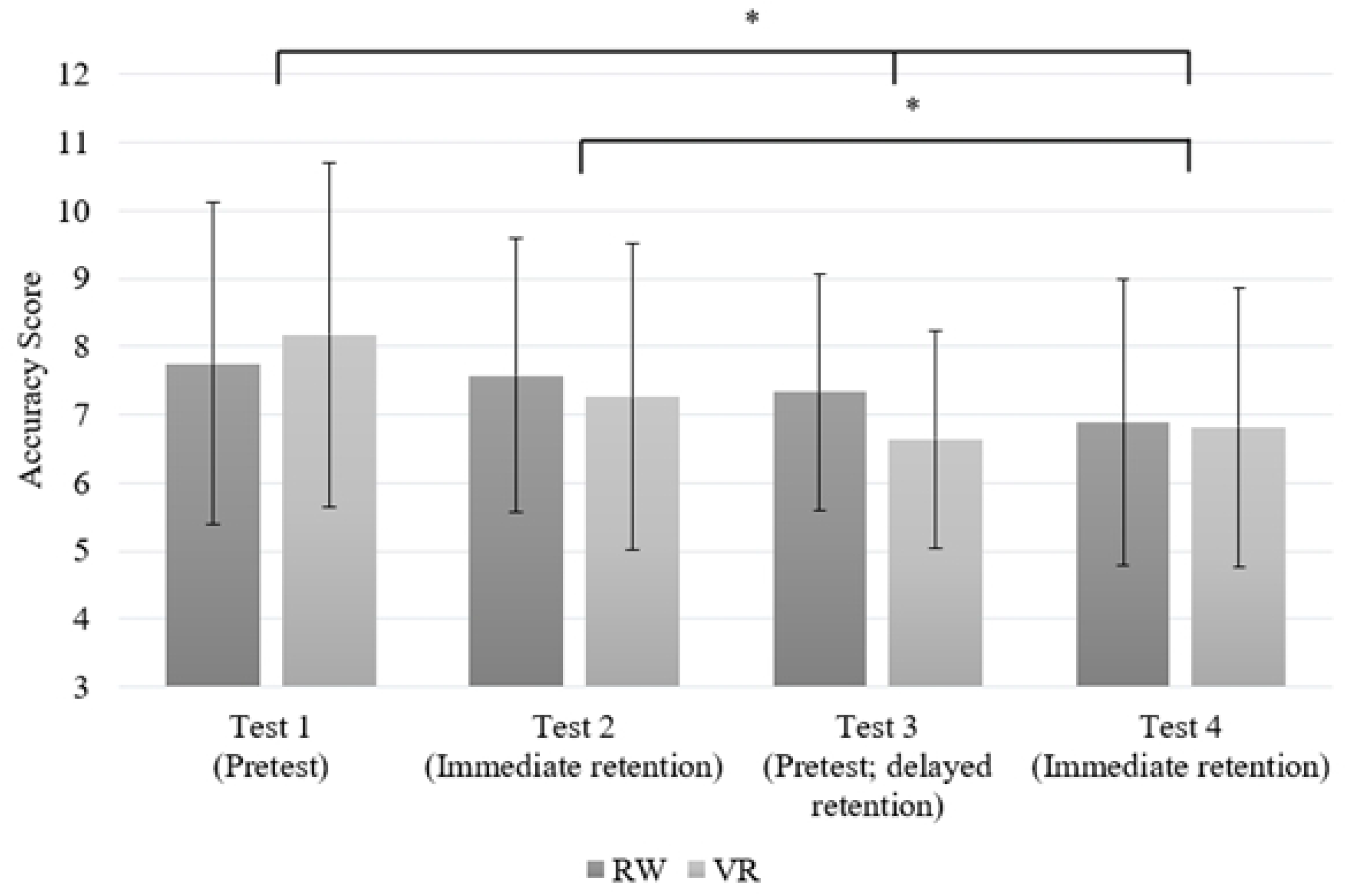
Accuracy score differences across tests. * p = < 0.05

### Kinematic Differences

#### Club Backswing

A 2 (group) x 4 (test) repeated measures analysis of variance (ANOVA) was used to determine kinematic differences during club backswing between groups and tests. Mauchly’s test indicated that the assumption of sphericity was violated (χ^2^(5) = 34.318, *p* < .001), therefore degrees of freedom were corrected using Greenhouse-Geisser estimates of sphericity (ɛ = .719). No significant differences were found between groups, *p* = .737, or within subjects across tests, *p* =.086.

#### Club Follow-through

A 2 (group) x 4 (test) repeated measures analysis of variance (ANOVA) was used to determine kinematic differences during club follow-through between groups and tests. Mauchly’s test indicated that the assumption of sphericity was violated (χ^2^(5) = 13.525, *p* < .019), therefore degrees of freedom were corrected using Greenhouse-Geisser estimates of sphericity (ɛ = .862). The analysis revealed a significant main effect for test *F*(2.587, 162.959) = 4.842, *p* = .005, *η_p_^2^* = .071. The interaction between test and group was not significant, *p* = .475. Additionally, the test of between-subject effects revealed a non-significant effect, *p* = .107. Given the significant main effect for test, pairwise comparisons were made to determine differences across tests (see figure 3). The analysis revealed that test two (*M* = .196, *SD* = .071) was significantly larger compared to test one (*M* = .176, *SD* = .063), *p* < .001. The analysis also revealed that test four (*M* = .191, *SD* = .067) was significantly larger than test one *(M* = .176, *SD* = .063), *p* = .006. Furthermore, comparisons showed that test three (*M* = .181, *SD* = .065) was significantly lower compared to test two (*M* = .196, *SD* = .071), *p* = .005.

**Figure 3.**
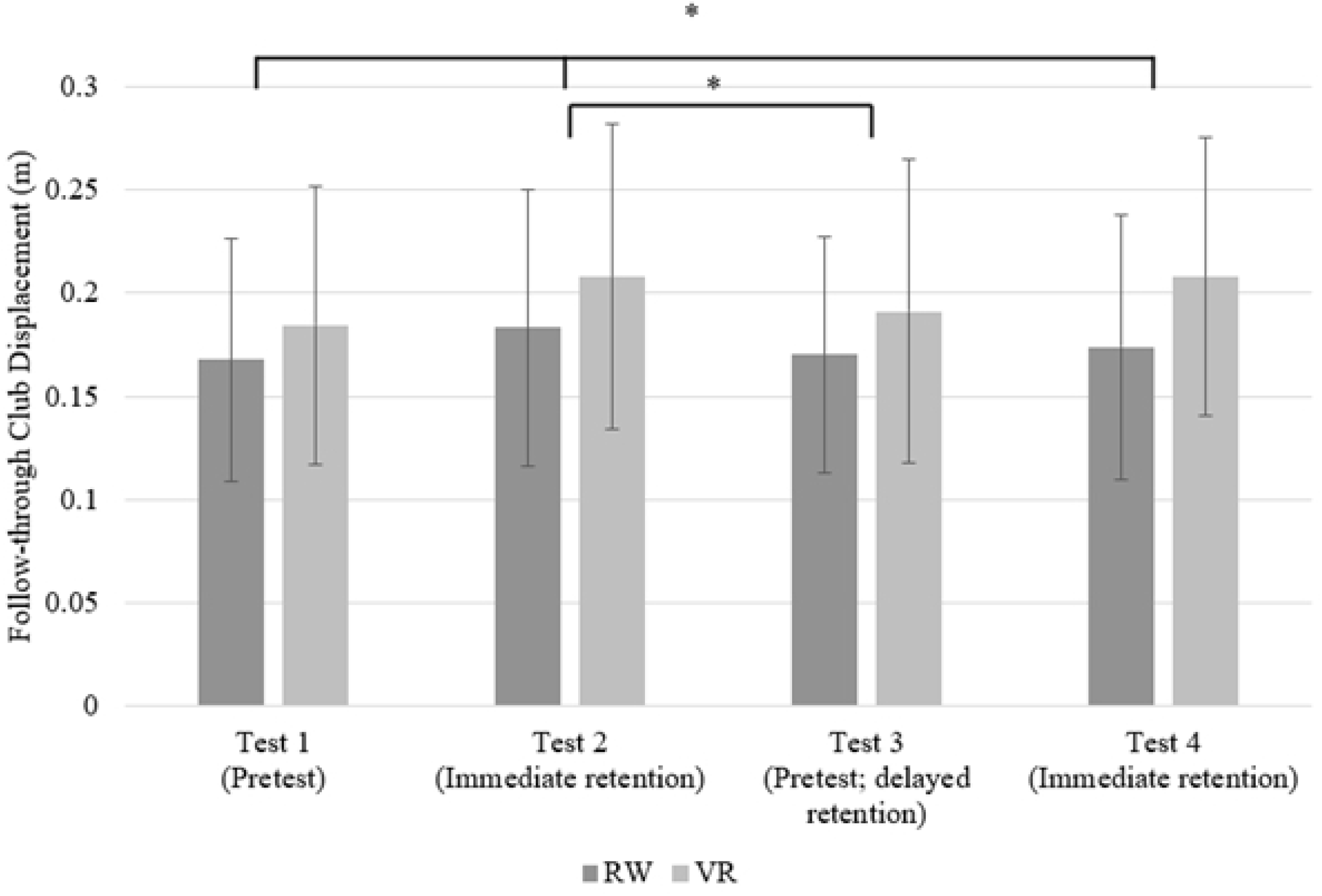
Follow-through displacement differences across tests. * p = < 0.05

#### Club Velocity

A 2 (condition) x 4 (test) repeated measures analysis of variance (ANOVA) was used to determine club velocity differences between groups and tests. The analysis revealed a significant main effect for test *F*(3, 183) = 2.682, *p* = .048, *η_p_^2^* = .042. No significant interaction between test and group was found, *p* = .735. Additionally, the test of between-subject effects revealed a non-significant effect, *p* = .051. Considering the significant main effect, pairwise comparisons for test were made to determine differences across tests (see figure 4). The comparisons revealed test three (*M* = .511, *SD* = .055) was significantly lower compared to test one (*M* = .524, *SD* = .071), *p* = .048 and test two (*M* = .527, *SD* = .057), *p* = .012.

**Figure 4.**
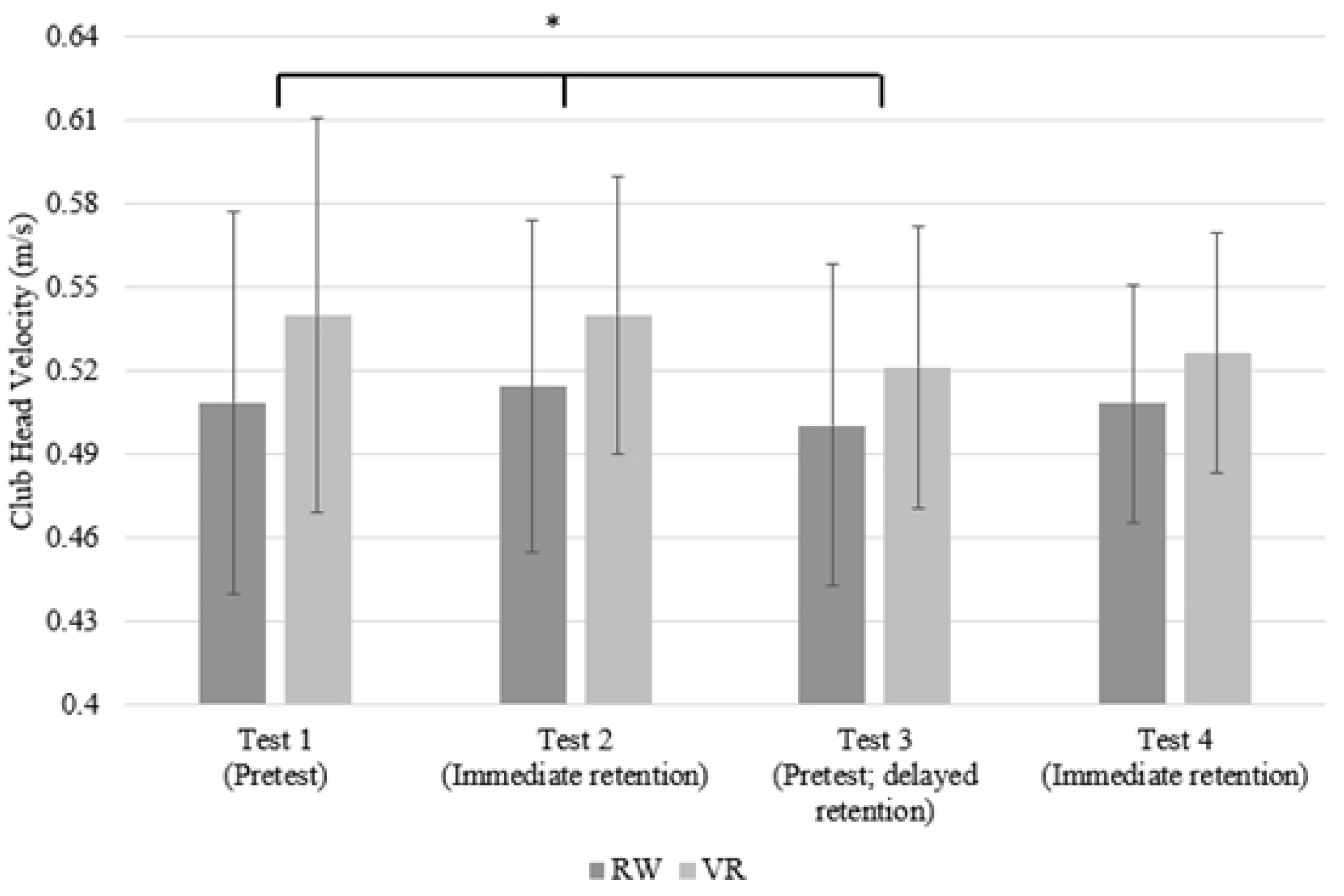
Club head velocity differences across tests. * p = < 0.05

## Discussion

The purpose of this study was to investigate the real-world golf putting accuracy and kinematic differences between RW practice and VR practice using a commercially available device and application. Based on previous research demonstrating positive transfer of learning following VR practice (9) and established theoretical frameworks (13,14,15), we predicted that both groups would improve motor performance, but no differences would be observed between groups. Based on Drew et al.’s (7) findings, we also hypothesized that there would not be group differences across the club swing kinematic measures. As predicted, both groups significantly improved golf putting accuracy at similar rates, and no differences were observed between the groups. Additionally, in line with our predictions, the analyses of the kinematic data revealed that the practice of putting a golf ball in VR or the real-world resulted in similar changes in real-world golf swing kinematics. In other words, swinging a virtual club to hit a virtual ball towards a virtual target resulted in putting accuracy and club swing technique improvements relative to actually swinging a real golf club to hit a real golf ball towards a real target. These results indicate that practicing a motor skill using commercially available VR technology can lead to similar RW performance improvements and biomechanical similarities when compared to RW practice.

The present study adds to the small body of research examining motor skill transfer within immersive VR. More so, this is only one of a few studies that have investigated the efficacy of using commercially available VR systems to improve real-world motor learning. Michalski et al. (9) was one of the first studies to establish that commercially available VR training is beneficial compared to no training at all. Here, and in line with Michalski et al. (9), this study demonstrates that practicing with commercially available VR systems can improve performance and appear to be equally effective compared to RW practice. Furthermore, these results are also in line with previous studies that have investigated the transfer of learning for both immersive (8 experiment 2,9,10) and non-immersive VR technology (2,17,18,19).

It is important to note that our findings are inconsistent with a recent study conducted by Drew et al. (7), in which they found that dart-throwing practice using an HTC Vive, hand-held controllers, and a commercially available application led to a decrease in real-world dart throwing performance. This negative transfer of learning likely occurred due to differences between the task performed in VR compared to the RW. Specifically, the virtual dart board height was scaled to the participant’s height in the virtual environment. In contrast, the dart board height in the physical environment was standard at a fixed height across all participants regardless of the height of the person. These task and environment differences reduced the similarities between conditions and likely led to the observed decrease in performance (12,13,16). In the present study, we scaled the virtual environment so the golf putting holes and putting distances were similar between the VR and RW conditions. Such similarities likely contributed to our observed positive transfer of learning.

In the present experiment, practice in both groups led to similar biomechanical measurements. Specifically, there were no club head displacement or velocity group differences in the posttest following VR or RW practice. While very few studies have examined performance production measures, such as biomechanical kinematics (1), a positive transfer within biomechanical measurements has been observed in both non-immersive (17) and immersive VR (7). The present results revealed that both groups similarly increased club head follow-through displacement and decreased club head velocity as a result of practice. A longer follow-through has been shown to be a characteristic of skilled putters (20,21). Therefore, the increased follow-through displacement observed in our study could have been due to an increase in skill level following the practice session. Regarding a decrease in velocity, the speed-accuracy tradeoff (i.e., Fitts’ law) suggests that individuals tend to exchange speed to maintain or increase levels of accuracy (22). Thus, is it not surprising to have observed a decrease in club velocity as individuals increased golf putting accuracy. Other golf putting research has indicated similar findings that a lower club head velocity is optimal for higher levels of accuracy (23,24).

To our knowledge, this is one of the first studies to show that practicing a motor skill using commercially available VR leads to similar results compared to physical practice. The use of a readily available golf putting application and hand-held controllers produced similar biomechanical characteristics and accuracy compared to RW golf putting. These results suggest that there might be value in using VR to supplement RW practice. However, given that other research has found a decrease in dart throwing performance following VR practice (7), understanding the generalizability of positive transfer of learning from commercially available VR is crucial, and the use of VR as a form of practice should not be used without prior transfer of learning validation. Future research should continue to investigate how the practice of other types of motor skills transfer to the RW environment in comparison to RW practice. Additionally, studies should investigate the extent to which the level of fidelity between virtual and physical realities can differ while still achieving a positive transfer of learning effect.

